# A statistical framework for mapping risk genes from *de novo* mutations in whole-genome sequencing studies

**DOI:** 10.1101/077578

**Authors:** Yuwen Liu, Yanyu Liang, A. Ercument Cicek, Zhongshan Li, Jinchen Li, Rebecca Muhle, Martina Krenzer, Yue Mei, Yan Wang, Nicholas Knoblauch, Jean Morrison, Siming Zhao, Yi Jiang, Evan Geller, Iuliana Ionita-Laza, Jinyu Wu, Kun Xia, James Noonan, Zhong Sheng Sun, Xin He

## Abstract

Analysis of *de novo* mutations (DNMs) from sequencing data of nuclear families has identified risk genes for many complex diseases, including multiple neurodevelopmental and psychiatric disorders. Most of these efforts have focused on mutations in protein-coding sequences. Evidence from genome-wide association studies (GWAS) strongly suggests that variants important to human diseases often lie in non-coding regions. Extending DNM-based approaches to non-coding sequences is, however, challenging because the functional significance of non-coding mutations is difficult to predict. We propose a new statistical framework for analyzing DNMs from whole-genome sequencing (WGS) data. This method, TADA-Annotations (TADA-A), is a major advance of the TADA method we developed earlier for DNM analysis in coding regions. TADA-A is able to incorporate many functional annotations such as conservation and enhancer marks, learn from data which annotations are informative of pathogenic mutations and combine both coding and non-coding mutations at the gene level to detect risk genes. It also supports meta-analysis of multiple DNM studies, while adjusting for study-specific technical effects. We applied TADA-A to WGS data of ∼300 autism family trios across five studies, and discovered several new autism risk genes. The software is freely available for all research uses.

## Introduction

*De novo* mutations (DNMs) arise spontaneously in offspring, and are often detected by sequencing families with disease occurrences, usually trios of parents and affected children (“trio-sequencing”). Researchers can identify risk genes using DNMs by searching for genes that harbor more *de novo* mutations in affected offspring than expected by chance. This approach has been highly successful in studying a range of developmental and psychiatric disorders including autism, intellectual disability, schizophrenia, epilepsy and congenital heart disease.^1–6^ DNMs tend to have larger effects than standing variants because they have not yet been acted on by natural selection. The DNM approach may be particularly helpful for early-onset diseases because standing risk variants for these phenotypes are rare and hard to identify with GWAS.

Most existing work on DNMs focuses on mutations in protein-coding regions. Even when whole genome sequence (WGS) data are available, researchers often analyze only the coding portion of the genome due to a lack of analytic tools for non-coding mutations.^7–9^ However, the majority of disease-associated variants identified by GWAS are located in non-coding sequences, potentially affecting gene regulation rather than protein function. This suggests that non-coding DNMs represent a large, currently unexplored, source of genetic variation that can aid gene discovery. The knowledge of non-coding disease variants will provide additional benefits. As the activity of regulatory elements tend to be cell type-specific, the analysis of DNM could offer clues as to which cell types are most relevant to disease etiology. A key research challenge is thus to provide an analytic framework for DNM data that incorporates non-coding mutations from WGS studies of disease families.

Current tools for DNM analysis perform some kind of “burden test” which evaluates whether the number of mutations in a gene is larger than would be expected by chance. He et.al. (2013) propose the method TADA for DNM analysis which effectively performs a weighted Bayesian burden analysis.^10^ TADA divides all mutations into categories, such as nonsense and missense mutations. Mutations in each category are weighted according to how damaging they are expected to be, with the weights for each category learned from the data. Another method, FitDNM, similarly performs weighted burden analysis, however, the weights are assumed to be known (from external source) instead of being estimated from data.^11^

Unlike protein-coding sequences, there is no simple genetic code for researchers to predict the functional effects of non-coding mutations, and thus difficult to assign them to simple categories. Instead, we can describe non-coding mutations using a number of overlapping genomic annotations such as tissue-specific epigenomic marks and cross-species conservation. Which annotations are relevant to a particular disease is not known *a priori*. Additionally, each annotation may only be weakly informative of pathogenic variants so we may need to combine multiple annotations. Existing tests developed for *de novo* coding mutations can not handle such complications: TADA can only handle disjoint categories of DNMs and has only been used with a small number of mutational categories; FitDNM is designed for exome sequencing data, assuming that the probability of a variant affecting protein function is known (from PolyPhen2).^11^

In this work, we present a statistical framework for analysis of DNMs, which we call TADA-Annotations (or TADA-A). TADA-A uses a probabilistic model of mutation counts for each position in the genome (Figure 1A). Specifically, we model the mutation counts as following Poisson distribution, and the background mutation rates depend on covariates such as types of nucleotide changes and local GC content. We expect the mutation rates in positions assigned to a disease-associated gene to be elevated compared with background rates, and the fold increases depend on functional annotations in a log-linear model. TADA-A offers several features important for WGS-based DNM studies. First, the model can take an arbitrary number of possibly overlapping annotations, and learn from the data which annotations are enriched for causal mutations. No arbitrary weighting scheme or variant filtering is needed. Secondly, the method predicts risk genes by combining information in both coding and non-coding regions. In addition, the information from coding mutations may come from an independent study, allowing a WGS study to borrow strength from published Whole Exome Sequencing (WES) studies. Finally, TADA-A supports meta-analysis of multiple WGS studies. It adjusts for possible difference in technical factors across studies by fitting a different background mutation model for each study.

**Figure 1.**
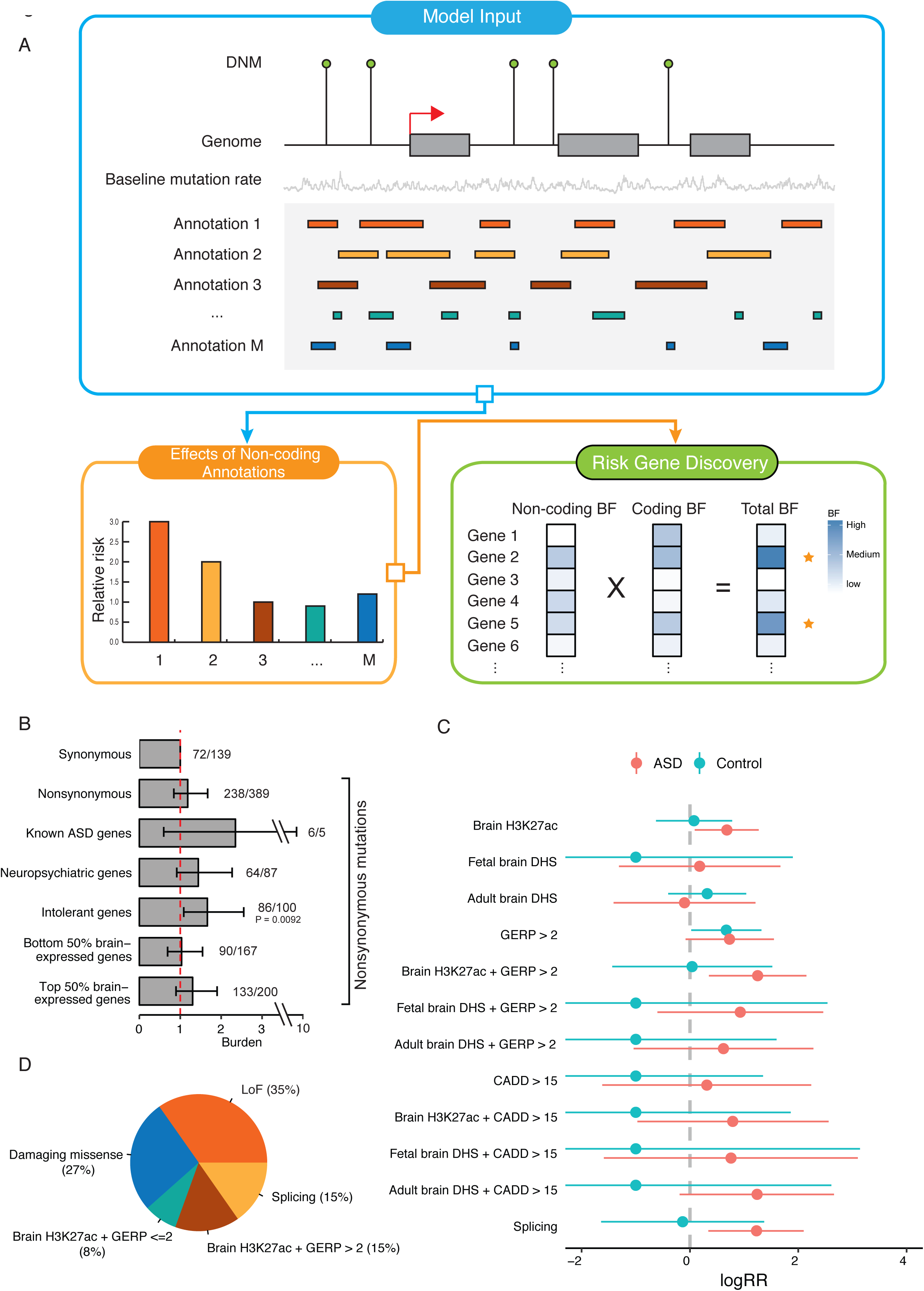
A. Overview of TADA-A. The blue frame illustrates the inputs of the model, including mutation counts, baseline mutation rates and annotations (assumed to be binary). The orange frame shows an example of relative risk estimates of different noncoding annotations by TADA-A The green frame illustrates our gene mapping strategy. For each gene, we derived its noncoding BF based on the relative risks of its noncoding mutations and calibrated mutations rates, which is then multiplied to the gene’s coding BF to get a total BF. B. Burdens analyses of different types of *de novo* nonsynonymus mutations. The error bars represent the 95% confidence intervals of burdens (ORs), based on Fisher’s exact tests. On the top of each bar, we labeled the number of mutations in ASD followed by in control. C. Estimated relative risks of different annotations using ASD DNMs and control DNMs. The x-axis is the Log(Relative risks). The error bars represent the 95% confidence intervals. D. Partition of *de novo* ASD risk into coding and non-coding mutations.

We apply TADA-A to study the contribution of non-coding sequences in autism spectrum disorder (ASD). WES studies using DNMs in autistic families have identified more than 70 ASD risk genes, highlighting the importance of DNMs in the study of autism.^1, 12–14^ Recently, efforts have been expanded to whole genome sequencing of ASD families. Two studies reported modest enrichment of functional non-coding DNMs near known ASD genes in autistic children, comparing with controls or unaffected siblings.^8, 15^ However, none of the published work has utilized non-coding DNMs to map specific risk genes or functional elements. We use TADA-A to analyze a collection of five whole-genome DNM datasets, leveraging a number of genomic annotations. We find that brain enhancers marked by H3K27ac, conserved brain enhancers marked by H3K27ac and high GERP scores (GERP >2), and regions predicted to affect splicing have increased rates of DNMs in ASD cases. Our conservative estimates suggest that regulatory non-coding mutations contribute to about a third of *de novo* autism risk (i.e. autism risk attributable to all DNMs). Using the DNMs from WGS data as well a published WES study, we were able to identify four new ASD risk genes at a FDR < 0.1. Multiple lines of evidence support the possible roles of these genes in ASD.

## Methods

### TADA-A model

TADA-A works in two stages: first, it calibrates the background mutation rates (mutation model); second, it learns which functional annotations are predictive of causal mutations and infer the risk genes (functional model). In the mutation model step, we assume that we have un-calibrated base-level mutation rates (summation over all possible allele-specific mutation rates at each base) from external data, e.g. from human-chimp comparison. We used the trinucleotide-based mutation rates table from Samocha et al. as our baseline rates.^16^ These baseline mutation rates are solely based the intergenic divergence between humans and chimps. For a particular study, observed mutation rates may differ from these un-calibrated rates as a result of study-specific technical factors. For example, lower sequencing depth reduces the number of called DNMs. Mutation rates may also depend on local genomic features such as GC content. To account for this variability, we calibrate the background mutation rates for each study. To simplify the computation, TADA-A collapses DNMs in a 50-bp genomic window into a single count. We model these mutation counts as Poisson Generalized Linear Model (GLM):

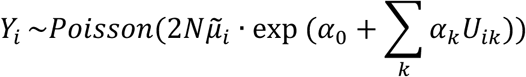

where *N* is the number of individuals, 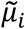 is the un-calibrated baseline mutation rate of window *i* (summing up the mutation rates of bases in window *i*), and the exponential term represents the deviation of the actual mutation rate of window *i* from 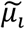. The variable *U_ik_* is the *k*-th mutation-related feature of window *i* and *α_k_* represents the effect of mutation feature *k*. The mutation features may include GC content, whether a sequence is transcribed, etc. TADA-A uses glm() in R to estimate the coefficients of the genomic features. After fitting the model, the calibrated allele-specific mutation rate at each base is the un-calibrated allele-specific mutation rate multiplied by a factor, exp(*α*_0_ + Ʃ_k_*α*_k_*U_ik_*) which is calculated for the window containing that base.

In the functional model, we model the dependency of allele-specific mutation rates on the gene status (risk gene or not) and functional genomic annotations. We make the functional model allele-aware because many annotations are allele-specific. For example, a *de novo* SNV could be nonsynonymous or synonymous, depending on what the mutant allele is. Noncoding annotations, such as CADD and SPIDEX scores are also dependent on the genotypes of mutant alleles. As described in the main text, we assume all mutations have been uniquely assigned to genes. Let *z_g_* be a binary indicator of whether gene g is a risk gene or not. When gene *g* is a risk gene (*z_g_* = 1), the number of DNMs mutating to allele *t* at base *i* aggregated from affected individuals follows a Poisson distribution

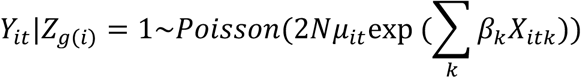

where *g*(*i*) is the gene that base *i* belongs to, *μ_it_* is the calibrated mutation rate to allele *t* of base *i* from the previous step, *X_it_* is the *k*-th genomic annotation of base *i* if mutated to allele *t*, and *β_k_* is the effect of the *k*-th annotation. Note that we consider annotations related to function at this step, such as conservation and enhancer activity. Since the annotations are binary, *e*^βk^ is the relative risk of the *k*-th annotation, i.e. the fold increase of mutation rates in positions with that annotation vs. those without the annotation. If gene g is a non-risk gene (*z_g_*= 0), the number of DNMs mutating to allele *t* at base *i* simply follows 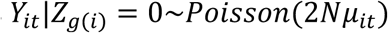.

Let *Y_g_* be the number of DNMs assigned to gene g. Assuming that DNM events are independent, the likelihood of *Y_g_*; given *Z_g_*, *P*(*Y_g_*|*Z_g_*), is simply the product of the probabilities of DNM counts at all bases over all possible mutant alleles, according to the equations above. Let *π_g_* be the prior probability of gene *g* being a risk gene, we have the full likelihood over all genes:

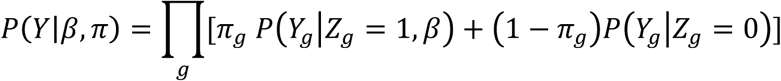

TADA-A implements two options for estimating parameters, both based on maximum likelihood. In the first option, *π_g_*; is the same for every gene, and we estimate its value by maximum likelihood jointly with *β*. In the second option, we use informative priors for *π_g_*; of all genes from external data, and we only estimate *β*. The confidence intervals of the parameters are based on standard asymptotic approximations using Fisher information matrix. When we have multiple annotations in the model, TADA-A uses a standard feature selection protocol to choose annotations. Specifically, it first fits a model with each single annotation and selects those whose coefficients are significantly different from 0 (at 95% confidence interval). It then refits the model jointly with the selected features.

Once we estimate the parameters, we compute the Bayes factor (BF) of a gene, as

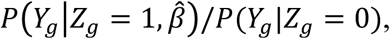

where the probabilities are evaluated at the MLE of parameter values. In our ASD analysis, we further multiply the BFs from non-coding analysis with the BFs from previous results based on coding mutations to obtain final BFs for all genes. We control for multiple testing using the Bayesian FDR control procedure.^17^

Because our likelihood is defined over all bases and all possible mutant alleles, including those possessing no DNM events, naïve parameter estimation is computationally expensive. To alleviate this computational burden, we reformulate the likelihood function by collapsing mutations over all bases sharing the same set of annotations, assuming all annotations are discrete. This strategy greatly reduces the computation time (Supplemental Methods). TADA-A software is available at GitHub (Web Resources).

### DNMs from whole genome sequencing data

The detailed information for each WGS dataset is summarized in Table S1. To remove erroneously called *de novo* SNVs, we excluded 8 individuals with more than 140 (2 times more than the median of ASD DNMs per individual) DNMs and removed all recurrent DNMs (i.e. exactly the same mutation in multiple individuals). Our unpublished (Wu et al.) DNM data are from WGS of 32 ASD trios of Han Chinese ancestry (Accession number: EMBL-EBI: PRJEB14713, data URL provided in Web Resources, details in Supplemental Methods,). These filtered data were used for all the analyses in this manuscript.

We also tried filtering out DNMs with a high allele frequency in GnomAD or BRAVO databases, as this could be one way of filtering sequencing errors. There are 167 mutations in cases and 18 in controls that have allele frequency more than 0.01 in either GnomAD or BRAVO. These mutations are not found in any of the novel ASD genes we identified. We found that removing these mutations did not change the model parameters (Figure S4), hence it will have little impact on our results.

### Non-coding annotations used in analyzing ASD data

Histone modifications: We used H3K27ac sites in fetal and adult brains to define cis-regulatory regions. Fetal brain sites from human cortex at embryonic stages 5, 7 and 12 p.c.w. were obtained from a recent study.^18^ For each stage, only peak regions consistent between two biological replicates were selected. Adult brain H3K27ac sites were obtained from Roadmap Epigenomics Project.^19^ They include regions from human angular gyrus, anterior caudate, cingulate gyrus, middle hippocampus, inferior temporal lobe, mid-frontal lobe and substantia nigra. We used MACS2 to call peaks from raw data, and only kept peak regions consistent between two biological replicates for each brain region. We used BEDtools^20^ to merge H3K27ac sites from fetal and adult brain.

DNase I hypersensitivity sites: Fetal brain DNase I sites were downloaded from Roadmap Epigenomics (male and female fetal brain) and Adult brain DHS data downloaded from ENCODE (Cerebrum_frontal_OC, Frontal_cortex_OC and Cerebellum_ OC).

Conservation scores: We used ANNOVAR to obtain GERP++ scores for all mutations.^21, 22^ We binarized GERP (a base is considered to be conserved if GERP is greater than 2).

CADD: We downloaded publicly available CADD scores (default parameters, v1.3) and binarized the scores (deleterious if one allele has a CADD score greater than 15).

Splicing score: We used results from SPIDEX, a deep learning-based approach to annotate variants that may affect splicing.^23^ An SNV is considering a splicing SNV if its delta-psi score is less than -1.416, which is the 10^th^ percentile of all positions with SPIDEX scores.

### Meta-analysis strategy and Applying TADA-A to ASD

In the ASD study, we first calibrate mutation rates of each study separately using the mutation model of TADA-A (Poisson regression), as described in Results. We then fit the functional model of TADA-A with non-coding annotations listed in the previous section. Since the calibrated mutation rates for any base may differ between studies, we calculated the likelihood of all genes for each study separately, and then multiplied the likelihoods over all studies to get a total likelihood. Note that the coefficients of the functional annotations are shared among multiple studies. We then estimate parameters via maximum likelihood. We take advantage of a previous Autism study to set the prior probabilities of risk genes *π_g_*.^2^ Specifically, we convert the Bayes factors reported in that study to posterior probabilities (assuming each gene has 6% chance of being ASD gene), and use these probabilities as *π_g_*. For computational reasons, we used the top 1000 genes, ranked by *π_g_*, in estimating the annotation parameters, since these genes are the most informative of parameters. After the first round of feature selection using all twelve annotations, only brain H3K27ac, brain H3K27ac + GERP > 2, and splicing effects had significant effect sizes, so we refit the model with only these features jointly. In the feature selection step, we define the search space of log(relative risk) to be from -1 to 10 to cover a wide range of possible effect sizes. Some annotations have the log(relative risk) estimated as -1 due to this boundary limitation.

To validate our relative risk estimation results, we used two different sets of informational priors. The first set is based on the FDRs of predicted ASD genes using a human brain-specific interaction network.^24^ For each gene, we derived its prior as 1 – FDR (We used the top 1000 genes for estimating relative risks.). The second set is based on a collection of 2601 genes implicated in neuropsychiatric disorders (See the definition of “Neuropsychiatric genes” in the next section for details). We assigned each neuropsychiatric gene a prior 0.431 to make the expected number of ASD genes consistent with the estimate that 0.06 of 18665 protein coding genes are ASD risk genes.^10, 25^

To identify specific ASD risk genes, we first derived the noncoding BF of each gene using data from each WGS dataset, then multiplied these BFs to get a total noncoding BF for each gene. This BF for a gene is multiplied with the published coding BF based on WES studies^2^. We then estimated q-values as mentioned previously.^17^ We used *π* = 0.06 as the fraction of ASD risk genes in this step.^10, 25^ We call a gene “novel ASD gene” (at a particular q-value cutoff) if its final q-value falls below the cutoff and its coding q-value from the previous study is above the cutoff. A gene that has no noncoding DNM will not be considered a new finding (even if a gene has no evidence from non-coding mutations, its q-value could still change).

### Definition of gene lists used in the analysis

Known ASD genes (194 genes) include genes with q-value < 0.3 from a combined analysis of CNVs, indels, and WES data using TADA^2^, SFARI category I (high confidence) and II (strong candidate) genes. Neuropsychiatric genes (2601 genes) are a larger set of genes likely involved in neuropsychiatric disorders, including genes with TADA q-value < 0.5,^2^ SFARI genes (Category I high confidence, Category II strong evidence), AutismKB genes,^26^ ASD risk genes summarized in a previous study,^27^ intellectual disability genes,^28^ the union of gene sets enriched with SCZ *de novo* coding mutations,^29^, high confidence postsynaptic density genes^30^ and FMRP targets.^31^ The set of nonASD genes are the 1000 genes with the highest TADA q-values.^2^ Intolerant genes include genes with top 5% RVIS^32^ and haploinsufficient genes obtained from two sources, one using copy number variations (genes with predicted haploinsufficient probability greater than 0.95),^33^ and the other using estimated mutation rate.^34^ To define tolerant gene, we started with genes with RVIS scores in the bottom 10%,^32^ genes with haploinsufficient probability smaller than 0.1,^33^ and genes that were used as control genes for LoF deficient genes.^34^ We then removed any genes that were in the intolerant gene set. To define gene groups based on their expression levels, we used the average expression level for each gene across all developing brain tissues from BrainSpan data.

### Burden analysis of different types of *de novo* coding mutations

In our burden analysis, we accounted for the difference in mutation rates between ASD subjects (∼60/individual) and controls (∼39/individual) using “background sequences” (sequences/mutations not expected to have function). Specifically, to assess the burden of nonsynonymous DNMs in ASD vs. controls, we used the numbers of synonymous SNVs in ASD and controls used as background.

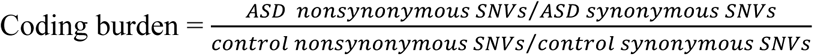

We tested if nonsynonymous DNMs were enriched in ASD vs. controls using Fisher’s exact test, and the burden was defined as the odds ratio (OR) from the 2 by 2 test.

### Assessing contribution of DNMs to ASD risk

We treated ASD liability (risk) as a continuous trait, and estimated the percentage of variance in ASD liability explained by four types of mutations. The variation of ASD liability explained by the *j-th* type of mutations is expressed as : 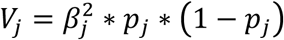 where *β_j_* is the effect size of the *j-th* type of mutations at the liability scale and *p_j_* is the probability that an individual carries a mutation of type *j* (see Supplemental Methods). Note that only causal mutations contribute to ASD liability, so both *β_j_* and *p_j_* are defined for mutations affecting causal genes. We calculated *β_j_* from the relative risk of j-th type of mutation using standard quantitative genetic calculations. To obtain *p_j_* we calculated the total mutation rate of type *j* mutations, and then multiply this by 0.06 (fraction of ASD risk genes) to obtain the rate of causal mutations of type j.

### Network analysis of candidate ASD genes

We used two tools, DAWN and GeneMania to analyze the connectivity pattern of our candidate ASD genes in gene networks. DAWN (Detecting Association With Networks) algorithm^35, 36^ is a guilt-by-association-based gene prediction algorithm. Its fundamental assumption is that risk genes tend to be functionally related with each other, and thus tightly connected in gene networks. A gene has a high posterior risk probability if it has a high prior risk probability, interacts in a network with other risk genes, or both. The prior risk probabilities came from published WES results.^25^ For the underlying network, we constructed partial co-expression networks for two spatial-temporal windows: the mid-fetal prefrontal cortex (PFC) and the infancy mediodorsal cerebellar cortex (MD-CBC), which are indicated as high risk windows for ASD.^37^ BrainSpan microarray dataset is used as the source for spatial-temporal gene expression data. DAWN was run separately for each above-mentioned network. We used regularization parameter (lambda) = 0.12, p-value cutoff = 0.1 and correlation thresholds 0.7 for PFC and 0.85 for MD-CBC, respectively. In Table 2, posterior risk scores (q-values) are shown for the candidate genes. A dash means that the corresponding gene is not co-expressed with other risk genes in any of the spatial-temporal windows.

**Table 2.**
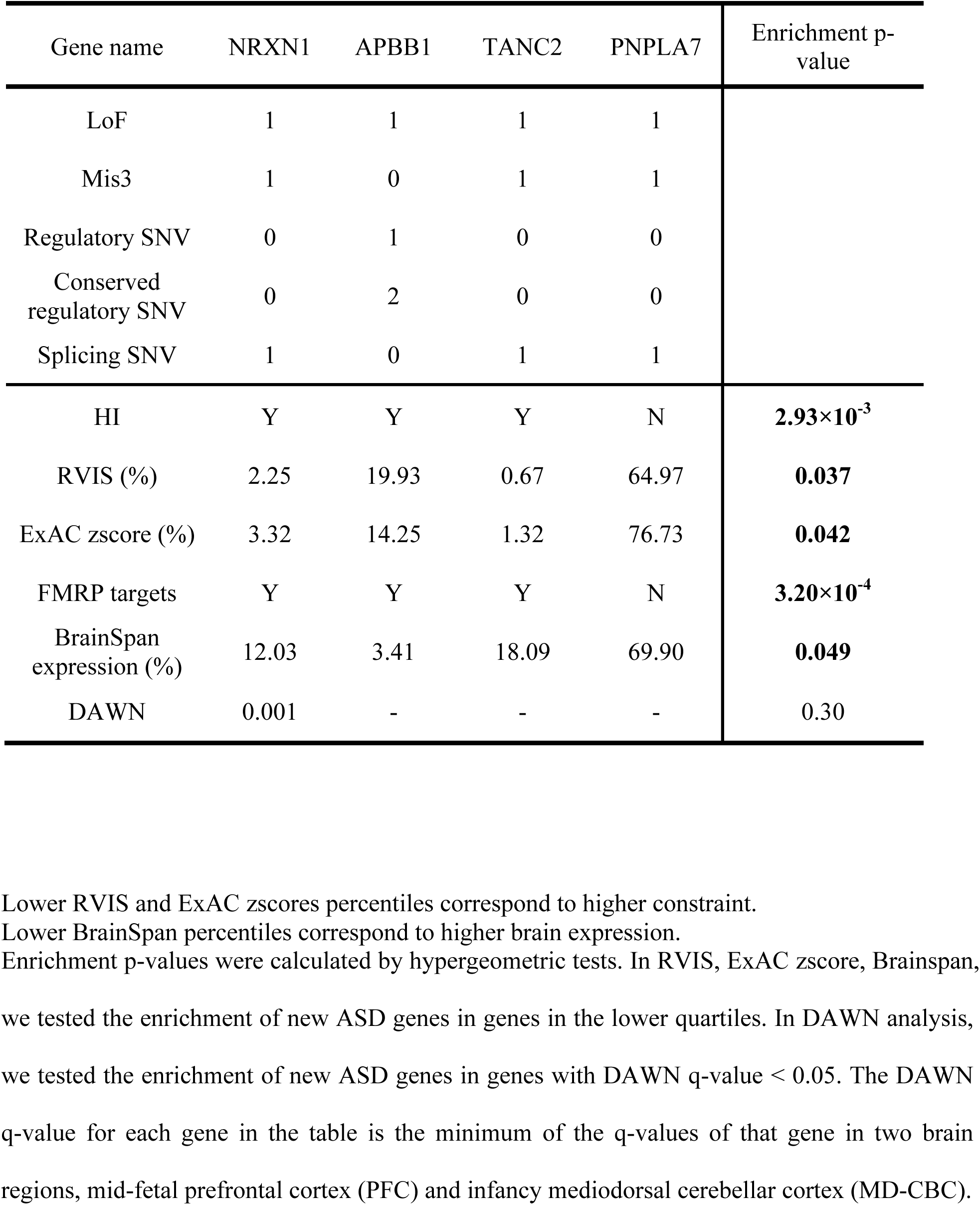
Mutational counts and evidence of four new ASD genes. In the evidence rows, Y means overlap with a gene set and N otherwise.

GeneMania^38^ is a tool for studying interactions among genes in a network using various types of information, such as gene co-expression and protein-protein interactions (PPIs). We studied the connection between our candidate genes with high-confidence ASD genes (genes with coding TADA FDR < 0.1 and genes in SFARI categories I and II) using co-expression data. The significance of the number of connections is assessed by randomly sampling gene sets of the same sizes as the candidate genes.

### Enhancers with recurrent *de novo* SNVs

We used all brain H3K27ac regions not overlapping with exons (not limited to sequences within 10kb of TSS). We observed 25 enhancers with at least two *de novo* SNVs in ASD samples, and we performed simulations to assess significance. In each simulation, we randomly re-distributed *de novo* SNVs of all brain enhancers, following a multinomial distribution. The multinomial probability of an enhancer is the ratio between the calibrated mutation rate of that enhancer and the sum of calibrated mutation rates across all enhancers (For each study, we first calibrated the trinucleotide-based mutation rates of all enhancers, accounting for sample size, GC contents and local 1Mb human-macaque divergence. We then added up this study-specific calibrated mutation rates across the five studies to get the total calibrated mutation rates for each enhancer). We performed simulations 10,000 times and obtained the distribution of the number of enhancers with recurrent SNVs.

### Power analysis

We generated *de novo* mutation data for all genes in the human genome (∼18700 genes) using the TADA-A model (See Supplementary methods for details). Briefly, we performed 5 simulations for each sample size, defined as the number of trios. For each iteration, we randomly assigned genes to ASD risk genes with a probability of 0.06, based on previous estimates.^10, 25^ For each risk gene, we sampled DNMs from each category (LoF, Mis3, less conserved regulatory SNVs, conserved regulatory SNVs, and splicing SNVs) according to allele-specific mutation rates and the average relative risks of these mutational categories, based on the TADA-A model. For non-risk genes, we set the relative risk at 1 for all mutational categories. We then used TADA-A to assess the evidence for each gene, using either coding mutations (WES approach) or all types of mutations (WGS approach). We then identified ASD risk genes with q-values < 0.1. To study cost-effectiveness between WES and WGS, we translated trio sample size into budget (WES: $500/sample; WGS: $1000/sample), and compared the number of identified ASD risk genes at each budget level.

### TADs with recurrent *de novo* SNVs

For each TAD region, we calculated the regulatory mutation rate as the sum of per-base calibrated mutation rates of brain H3K27ac sites within the TAD (For each study, we first calibrated mutations rates of these H3K27ac sites and their 2.5kb flanking regions, accounting for sample size, GC contents and 1Mb human-macaque divergence. We then added up the calibrated mutation rates across five studies together). Under the null hypothesis, the count of regulatory SNVs follows a Poisson distribution, whose rate is the regulatory mutation rate.^10, 16^ We then calculated the *p-*value of each TAD region using the Poisson test and used the Benjamini-Hochberg procedure to control FDR.

## Results

### Overview of TADA-Annotations (TADA-A)

TADA-A consists of two steps. In the first step, study-specific, background mutation rates are estimated at each position. We use an initial estimate solely based on a trinucleotide mutation rates table from the literature.^16^ These mutation rates are derived from the divergence in intergenic regions between humans and chimps, which are subject to less natural selection comparing with coding regions. We then adjust for genomic and technical covariates such as sequencing depth, local GC content, and 1-Mb local divergence scores between humans and macaques. In the second and main step, we use DNMs and annotation information to identify risk genes. We assume that we can assign each DNM to one gene, however, we could also analyze at the level of genomic regions and the DNM-to-gene assignment is not strictly necessary (see Discussions). TADA-A takes as input the number of DNMs at each genomic position, summing over all affected subjects, and a set of possibly overlapping, genomic annotations (Figure 1A, upper panel). Genomic annotations might include cell specific histone modifications or evolutionary conservation. TADA-A produces two main outputs: (1) the annotations that are informative of causal mutations and their effect sizes and (2) the predictions of specific susceptibility genes of the disease of interest (Figure 1A, bottom panel). We measure the effect of an annotation by its relative risk, i.e. the fold increase of disease risk for a variant carrying that annotation vs. a variant without that annotation (assuming the annotation is binary). The model of TADA-A is general enough that it can analyze either coding or non-coding mutations. If both types of mutations are analyzed, the results could be easily combined by multiplying the resulting Bayes factors (BFs).

The intuition behind TADA-A is that, in affected individuals, disease-causing mutations should appear at higher rates than expected from the baseline mutation rate. Our model can be written as 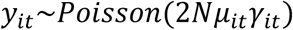, where *y_it_* is the observed number of *de novo* mutations at position *i* mutating to allele t (*y_it_* is usually 0 or 1), *μ_it_* is the expected background mutation rate estimated in the first step, N the sample size, and *γ_it_* is the relative risk of a DNM at position *i* mutating to allele t (greater than 1 for risk mutations). To model the relative risk, *γ_it_*, we define a binary (unobserved) variable for each gene indicating whether it is a risk gene or not. For a non-risk gene, all its positions have relative risk equal to 1. For risk genes, we model log (*γ_it_*) as a linear function of the genomic annotations. Each gene has a prior probability of being risk gene. TADA-A offers the option of using informative prior probabilities, e.g. a likely risk gene from previous WES studies would have high prior probability. Intuitively, this allows us to put more emphasis on highly plausible risk genes to estimate the parameters of annotations, while discounting the unlikely disease-associated genes. This is important when statistical signals in the annotations are weak.

We estimate model parameters (mainly the relative risk of each annotation) using maximum likelihood. Since annotations could be partially redundant (e.g. an enhancer may be associated with multiple annotations such as open chromatin and H3K27ac) and not all annotations are informative, we implement a feature selection protocol to first select annotations that are informational to predict pathogenic mutations and then jointly estimate the relative risks of these selected annotations. Once we have estimates of all the parameters, we predict whether a gene is a risk gene or not using Bayes factor (BF), combining information in all its associated DNMs. Similar to the original TADA method, we test each gene separately, and contrast the null model where the relative risk is always 1 with the alternative model described above for risk genes (relative risks dependent on annotations). To use TADA-A in a meta-analysis setting that combines multiple studies with possibly different rates and patterns of DNMs (e.g. in relation to GC content), we fit a different background mutation model for each study, but estimates a common set of parameters related to functional annotations.

TADA-A can be used to answer several questions about genetics of a complex disease, ASD in our case. What annotations are associated with causative mutations? Based on this knowledge, can we learn about the genetic architecture of the disease, especially about the relative contribution of coding vs. non-coding DNMs to the disease liability? Finally, can we identify specific disease-associated genes? We present below our results in answering these questions for ASD.

### Whole genome sequencing data of ASD and mutation rate calibration

We analyzed DNM data from five WGS studies of ASD trios or quartets, with a total of 314 affected subjects (Table S1). Mutation data is limited to *de novo* SNVs. The validation rate of *de novo* SNVs based on Sanger sequencing ranges from 85% to 94% in the five studies^39–41^. The number of DNMs per subject ranges from 57 to 63. Additionally, we collected the control data from a cohort of ∼700 non-ASD subjects. While TADA-A does not need control data, we use this additional dataset to perform burden analysis often employed in DNM studies, comparing the rate of DNMs in affected subjects with the rate in controls (see Methods). Our main results are limited to sequences close to genes, including protein-coding sequences, non-coding sequences within ±10 kb of TSSs, and potential splicing-regulatory regions that are not covered by these two categories. In a later section, we present results based on distal sequences.

To account for technical difference among studies that may affect observed mutation rates, we use published mutation rates from Samocha et al. as an initial estimate,^16^ and adjust for covariates using a Poisson regression model, separately for each of the five datasets (see Methods). We perform analysis at the level of 50bp non-overlapping sliding windows and consider four covariates for estimating baseline mutation rates: (1) whether a window is in coding regions (transcribed sequences may have lower mutation rates because of transcription-coupled repair); (2) whether the window is in promoter regions (CpG (de)methylation in promoters might affect mutation rates); (3) the percent GC in the window, which may correlate with sequencing depth and hence DNM detectability;^8^ (4) the divergence score between humans and macaques of the 1Mb window around the window, which is used to capture the local deviation from the trinucleotide-based mutation rates. The effects of these covariates are summarized in Table S2. In all of the five studies, the intercept in the regression model is significantly different from 0 *(P* < 0.05), suggesting systematic departure of average mutation rates from the published rates. In three of the five ASD studies, GC content has a negative effect on the observed DNM rate (*P* < 0.05 for three). The mutation features representing whether a sequence is in coding or promoter region were found to have a relatively large effect in specific studies (e.g. the coding feature in Jiang et al. was significant at *P* = 0.003). As expected, local divergence scores have a positive effect on the observed mutation rates in all of the five studies, though the effect is small and not statistically significant. The results from our mutation rate modeling thus support the importance of accounting for difference in studies in meta-analysis of DNM datasets.

We notice that although some studies have a small sample size, the numbers of DNMs are still much larger than the number of parameters. For example, even for the dataset with the smallest sample size (The data of Michaelson et al. has a sample size of 10), we still have 79 DNMs in the 50-bp windows that are included in our model. In addition, we fit mutation rate parameters separately for each study, so a study with small sample size will not impact the estimates for a larger study.

### Risk-increasing mutations in ASD are associated with active enhancer mark and damaging effects on splicing

We first assessed the quality of data using coding DNMs. We performed a simple burden analysis of protein-coding sequences in probands vs. controls. We adjusted for the difference in baseline mutation rates in the ASD studies and controls using synonymous mutations (whose true mutation rates should be the same across studies, see Methods). As expected, we found that the average rate of non-synonymous mutations per subject is about 1.2 fold higher in ASD subjects vs. controls (Figure 1B), in line with previous estimates.^2, 12, 42^ We also observed an increased rate of non-synonymous mutations in gene sets enriched with ASD risk genes, including known ASD genes, genes likely involved in neuropsychiatric disorders (dubbed “neuropsychiatric genes”), genes intolerant to mutations, and genes highly expressed in the brain (Figure 1B). Only the burden in mutation intolerant genes is statistically significant (*P* < 0.01). Synonymous mutations have recently been reported to be enriched in ASD cases as they may disrupt transcriptional regulatory processes, such as splicing.^8^ Thus we think that using synonymous mutations only makes our results more conservative: if there is indeed enrichment in synonymous mutations, the burden of nonsynonymous mutations would be under-estimated as a result of adjusting mutation rates using synonymous mutations.

For TADA-A analysis, we use a total of 12 functional annotations (Figure 1C). Some annotations measure the regulatory function of variants, including fetal and adult brain H3K27ac,^18, 19^ fetal^19^ and adult brain^43^ DNase Hypersensitive Sites (DHS). H3K27ac is a mark of active enhancers and DHS is a mark of open chromatin, often suggestive of regulatory functions. We also use a conservation score GERP,^22^ and an aggregate variant score CADD,^44^ and composite annotations of regulatory regions and GERP or CADD. Splicing has been shown to be important for many human diseases, so we include the splicing effects predicted by SPIDEX.^23^ We choose the 10^th^ percentile of SPIDEX scores as a cutoff. SNVs with a SPIDEX score smaller than this cutoff were classified as affecting splicing. These SNVs are enriched within 20 bp around exon/intron junctions (Figure S1).We limit our analysis to sequences within 10 kb of transcription start sites (TSSs) of protein-coding genes (not including UTRs) and potential splicing-regulating regions (which could be far away from TSSs). To increase the power of TADA-A to detect predictive annotations, we take advantage of existing WES studies. For each gene, we summarize the findings of previous WES studies as the probability of being an ASD risk gene^2^, which is then used as the prior probability of being a risk gene in the TADA-A model. This step allows us to put large weights on known ASD risk genes, whose probabilities are close to 1, comparing with average genes (about 0.06).

In our initial analysis of feature selection using TADA-A, we found that among 12 annotations, only brain H3K27ac, brain H3K27ac + GERP > 2, and the SPIDEX score, when estimated separately, make marginally significant contributions (p < 0.05, Figure 1C). We therefore retrain the model using only these three features and jointly estimate their relative risks at 1.54, 3.42, and 3.22 respectively (Table 1 and Table S3). In the following analysis, we refer to *de novo* SNVs in H3K27ac regions within 10kb of genes as regulatory SNVs (those with GERP > 2 as conserved regulatory SNVs), and *de novo* SNVs predicted by SPIDEX to affect splicing as splicing SNVs. To study if the splicing signal is robust, we also used another simple but commonly used way to predict splicing mutations. We predicted splicing SNVs as any SNVs that are within 20 bp windows of exon/intron junctions. The estimate of Log(Relative risk) is very close to SPIDEX prediction, though is less significant (logRR estimate is 1.09, lower bound is -0.18 and upper bound is 2.35). This difference may be due to the fact that many of the bases within 20bp of exon/intron junctions do not regulate splicing. We also tried using two other sets of informative priors to analyze the 12 annotations: one based on a genome-wide prediction of ASD genes in the context of a human brain-specific gene interaction network^24^ and the other based on a neuropsychiatric disorder gene set (see Methods for details). The resulting estimates for functional annotations are largely similar, suggesting that our estimates are quite robust (Figure S2A, Figure S2B).

**Table 1.** Mutation frequency (the number of mutations per subject) and the relative risks of different types of *de novo* mutations. Note that the mutation frequency and relative risk is based on mutations in risk genes.

| Mutation Class | Mutation frequency | Relative mutational exposure(%) | Relative risk | Variance of ASD liability explained (100%) |
| --- | --- | --- | --- | --- |
| Mis3 | 0.0175 | 12.5 | 4.70 | 0.83 |
| Loss-of-function | 0.00405 | 2.90 | 20.0 | 1.08 |
| H3K27ac SNV (Gerp <=2) | 0.0839 | 59.9 | 1.54 | 0.24 |
| H3K27ac SNV (Gerp > 2) | 0.0164 | 11.7 | 3.42 | 0.46 |
| Splicing SNV | 0.0183 | 13.1 | 3.22 | 0.46 |

In the analyses above, we borrowed priors from other studies to increase our power to detect non-coding signals which are generally weaker than coding signals. When using a uniform prior of 0.06 to perform relative risk estimation, we found that while the sign of several annotations, including brain H3K27ac + GERP > 2 and SPIDEX, remain the same (Figure S2C), the strength of statistical evidence is much weaker. This is consistent with our expectation and underscores the advantage of using informative priors to increase the sensitivity of signal detection.

To demonstrate that the signal discovered by TADA-A is specific to ASD, we run TADA-A using the control data. All the annotations now have relative risk estimates close to or smaller than 1, except for one feature (GERP > 2), though the lower bound of Log(Relative risk) is very close to 0 (0.024) (Figure 1C). In addition, combining this feature with other epigenomic features did not increase the effect size as it did when analyzing ASD data. So we believe that the result of this annotation is likely due to noise.

### Enhancer and splicing mutations make substantial contributions to the *de novo* risk of autism

A fundamental question in genetics is how the risk variants are distributed among various functional classes, such as protein-coding sequences, enhancer sequences, non-coding RNAs, etc. This question has been studied recently using common variants.^45, 46^ It was found that even though variants in protein coding regions are highly enriched with risk variants, they explain only a small fraction of total disease risk. The results from the previous TADA-A analysis allows us to address this “risk partition” problem from a different angle, using DNMs. Based on TADA-A results, we considered three types of non-coding *de novo* SNVs: regulatory SNVs with GERP <=2 (less conserved regulatory SNVs), regulatory SNVs with GERP > 2 (conserved regulatory SNVs) and splicing SNVs, in addition to two classes of coding mutations: LoF and probably damaging missense (predicted by PolyPhen-2, denoted as Mis3). We quantify the contribution of a mutation type as Liability Variance Explained (LVE), taking into account both the frequencies of this mutation type and its average relative risk (see Methods). For coding mutations (LoF and mis3), the relative risks were obtained from published TADA estimates in WES studies.^10^ For non-coding SNVs, we used the relative risks estimated by TADA-A.

The relative risks of regulatory SNVs and splicing SNVs are lower than those of coding SNVs (Table 1). Despite a lower risk per variant, regulatory SNVs are much more frequent than other classes of mutations, making the total contribution of regulatory SNVs comparable to LoF or missense coding mutations. Each class of mutation explains only a small fraction of estimated total ASD genetic risk (Table 1), consistent with the conclusion of an earlier study.^47^ Considering only the risk due to *de novo* mutations, we found that non-coding SNVs (including less conserved regulatory SNVs, conserved regulatory SNVs and splicing SNVs) explain 38% of the *de novo* risk (Figure 1D). This estimate, however, is likely very conservative (Discussion).

### TADA-A identifies novel ASD risk genes by combining coding and noncoding mutations

A recent WES study (∼3,500 samples) identified 58 ASD risk genes at FDR < 0.1^2^ using *de novo* SNVs. Applying TADA-A on the WGS data and combining them with the WES results, we discovered four new ASD risk genes at FDR < 0.1 (Table 2), and 12 at FDR < 0.3 (Table S4). Each of the four genes at FDR < 0.1 has at least one LoF or Mis3 mutation, and the evidence for these genes is strengthened by the presence of regulatory or splicing SNVs. We found extensive evidence supporting the plausibility of these genes as ASD risk genes. *APBB1*, *NRXN1, and TANC2* are the targets of neuronal-RNA binding protein FMRP, whose loss of function causes Fragile X syndrome and autistic features (Table 2 and Table S5, hypergeometric test, *P =* 0.00032). These three novel genes have been identified as haploinsufficient genes^33, 34^ (Table 2 and Table S5, hypergeometric test, *P =* 0.00293) and are highly expressed in the brain (Table 2 and Table S5, hypergeometric test, top 25% of all genes, *P* = 0.049). The genes also tend to be evolutionarily constrained as measured by either RIVS (Table 2 and Table S5, hypergeometric test, RVIS top 25% genes, *P* = 0.037) or another metric based on tolerance of LoF variants in ExAC (Table 2 and Table S5, hypergeometric test, ExAC top 25% genes, *P =* 0.042). Evolutionary constraint in the human population has been shown to be a strong predictor of autism genes^16^.

We performed network analyses to further establish the link of new candidate genes to autism. DAWN^35, 36^ is a recently developed method that predicts autism risk genes by virtue of the genes’ association with known ASD genes in co-expression networks of early developing brain. A gene receives a high DAWN score if it is highly connected with other likely ASD genes. We found that NRXN1 has a DAWN q-value < 0.05 in at least one of the two critical spatial-temporal developmental windows for ASD^37^ (Table 2 and Table S5, Fold of enrichment 2.92, though the hypergeometric test *P* = 0.30 is not significant, but the power of the test is small as there are only four novel genes). Using GeneMania,^38^ we found our candidate genes were highly connected to high-confidence ASD genes in the gene co-expression network constructed from multi-tissue gene expression data (91 co-expression links between the two gene sets*, P* = 0.04, Figure 2A). These analyses, using different analytic tools and genomic data, thus support that our new genes are functionally related to known ASD genes.

**Figure 2.**
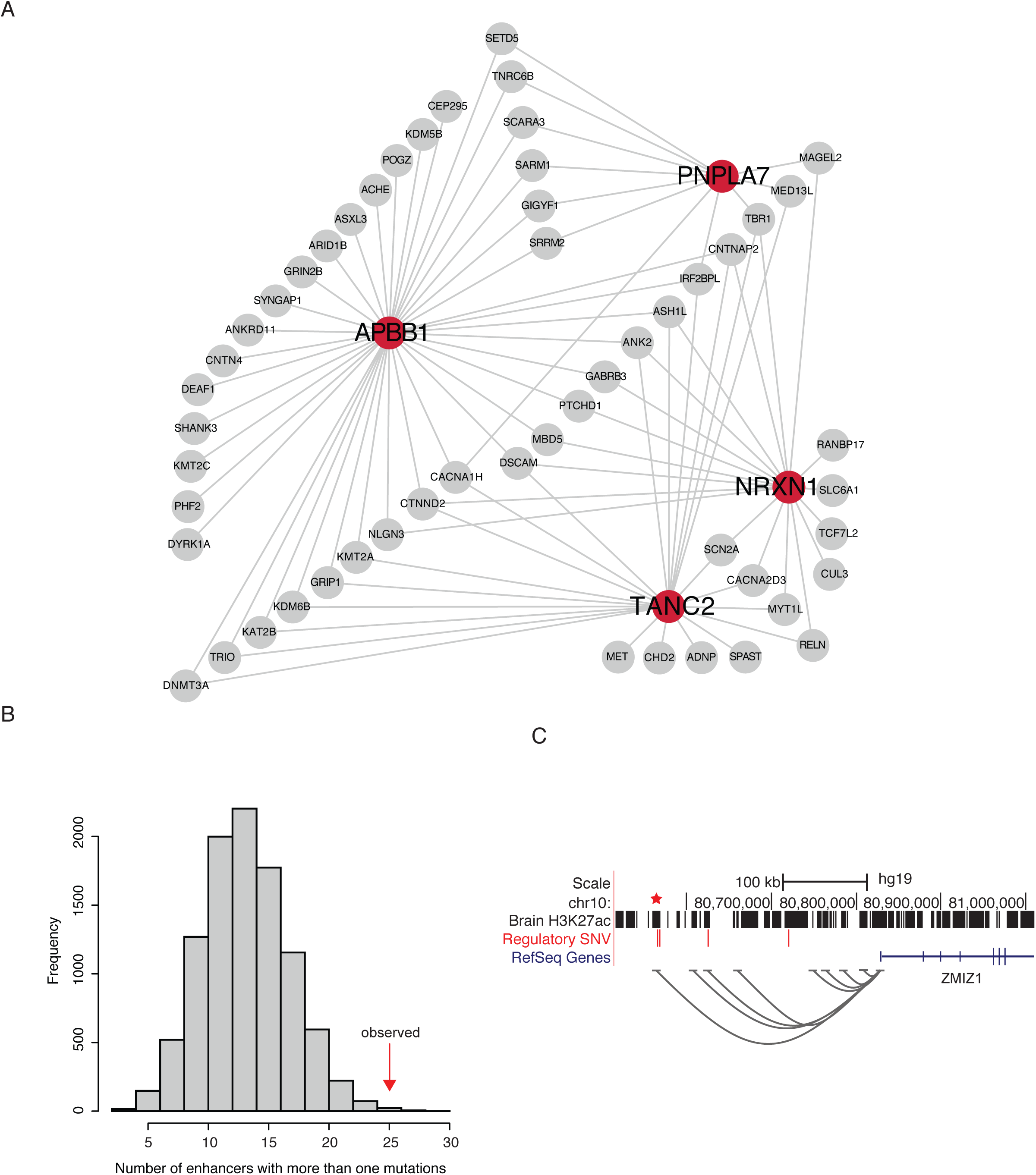
Predicted risk genes and enhancers of ASD. (A) GeneMania network analysis of the five new ASD genes. Red circles represent new ASD genes while grey circles represent known ones. Two genes are connected if their co-expression across multiple tissues reaches a threshold. Only connections between the two gene sets are shown (B) Distribution of the number of enhancers with recurrent (two or more) *de novo* SNVs from 10,000 simulations. The vertical red arrow marks the observed number of enhancers with recurrent *de novo* SNVs. (C) A distal enhancer (marked by a star) of ZMIZ1 with recurrent SNVs. Grey curves represent possible interactions between enhancers and promoters (correlated activities across multiple tissues). Note that the region contains two additional DNMs in other sequences.

Expanding our analysis to the 12 new genes at FDR < 0.3, we observed significant enrichment of multiple gene annotations (Table S4), including haploinsufficient genes (Table S5, hypergeometric test, *P =* 0.00035), constrained genes (Table S5, hypergeometric test*, P =* 0.0012 using RVIS and *P =* 0.0096 using variant frequency in ExAC), and genes significantly co-expressed with known ASD genes from DAWN analysis (Table S5, hypergeometric test*, P =* 0.0022). The 12 new genes are also significantly enriched in genes predicted to be ASD risk genes (FDR < 0.1) by a recently developed machine learning approach that utilizes a brain-specific functional gene network (Table S5, hypergeometric test*, P =* 0.028).^24^ Literature inspection provides further support of the roles of most of these genes in ASD (Table S6).

### Distal enhancers and TADs with multiple *de novo* mutations implicate additional risk genes

Our TADA-A analyses were performed at the gene level, and only considered enhancers within 10 kb of TSSs. Applying TADA-A to distal enhancers is challenging largely because of the uncertainty of assigning these enhancers to their target genes. Various studies have shown that only in 10-30% cases, distal enhancers target their nearest genes. We use a different approach in this section to test if distal enhancers may play some roles in autism. Our idea is that the probability of multiple DNMs occurring in a single enhancer by chance is very low. We found 25 H3K27ac enhancers with two or more SNVs in ASD cases, significantly higher than random expectation based on simulations (Figure 2B, *P =* 0.0014). We predicted the likely target genes of recurrent enhancers based on cross-tissue correlation between enhancer activity and gene expression from Roadmap Epigenomics (Table S7). We found a recurrent enhancer putatively targeting *ZMIZ1*, more than 250 kb away (Figure 2C). The region contains two other DNMs in two enhancers, one of which also has correlated activities with the ZIMZ1 promoter. A target of FMRP, *ZMIZ1* is highly expressed in the brain and interacts with neuron-specific chromatin remodeling complex (nBAF) which is important in regulating synaptic functions.^48, 49^ Several nBAF members have been linked to autism, such as ARID1B and BCL11A.^50^ The pathogenic potential of *ZMIZ1* is further supported by the observation of a *de novo* gene-disrupting translocation in an individual with intellectual disability.^51^ These results strongly support the role of ZMIZ1 in autism, and also highlight the mechanism that DNMs may increase ASD risk by disrupting distal regulatory elements.

We applied the similar idea of recurrent DNM analysis at the level of Topologically Associating Domains (TADs).^52^ These are megabase–sized chromatin interaction domains that are stable across cell types and have been proposed to demarcate transcriptional regulatory units.^52^ Based on estimated mutation rates, we found two TADs with a significant (at FDR < 0.3) number of regulatory SNVs (Figure S3 and Table S8). In both TADs, there are only two or three genes, and we conjecture that SRBD1 and MRSA are likely the underlying ASD genes in the two TADs, respectively (see Discussion).

### Power of mapping ASD risk genes with WGS and WES

Enlightened by the *de novo* genetic architecture of ASD (Table 1), we used simulations to address how the power of a DNM-focused WES or WGS study depends on its sample size and sequencing budget. We randomly sample ASD risk using a prior probability of 0.06 based on previous estimates of the total number of ASD risk genes^1, 53^; randomly sampled mutations according to mutation rates and the TADA-A model (causal genes tend to have more deleterious coding and non-coding mutations compared to expectations) and then applied TADA-A to identify risk genes at q-value < 0.1. We found that the power of the simulated WGS design is about 50%-120% higher than that of the WES (Figure 3A). The gain of power by WGS is more obvious when the sample size is smaller. We next investigated whether the additional power gained from WGS is justifiable on the basis of cost. At the current per-sample cost level (WES: $500 and WGS: $1000), we found that WES is still more cost-effective than WGS (Figure 3B).

**Figure 3.**
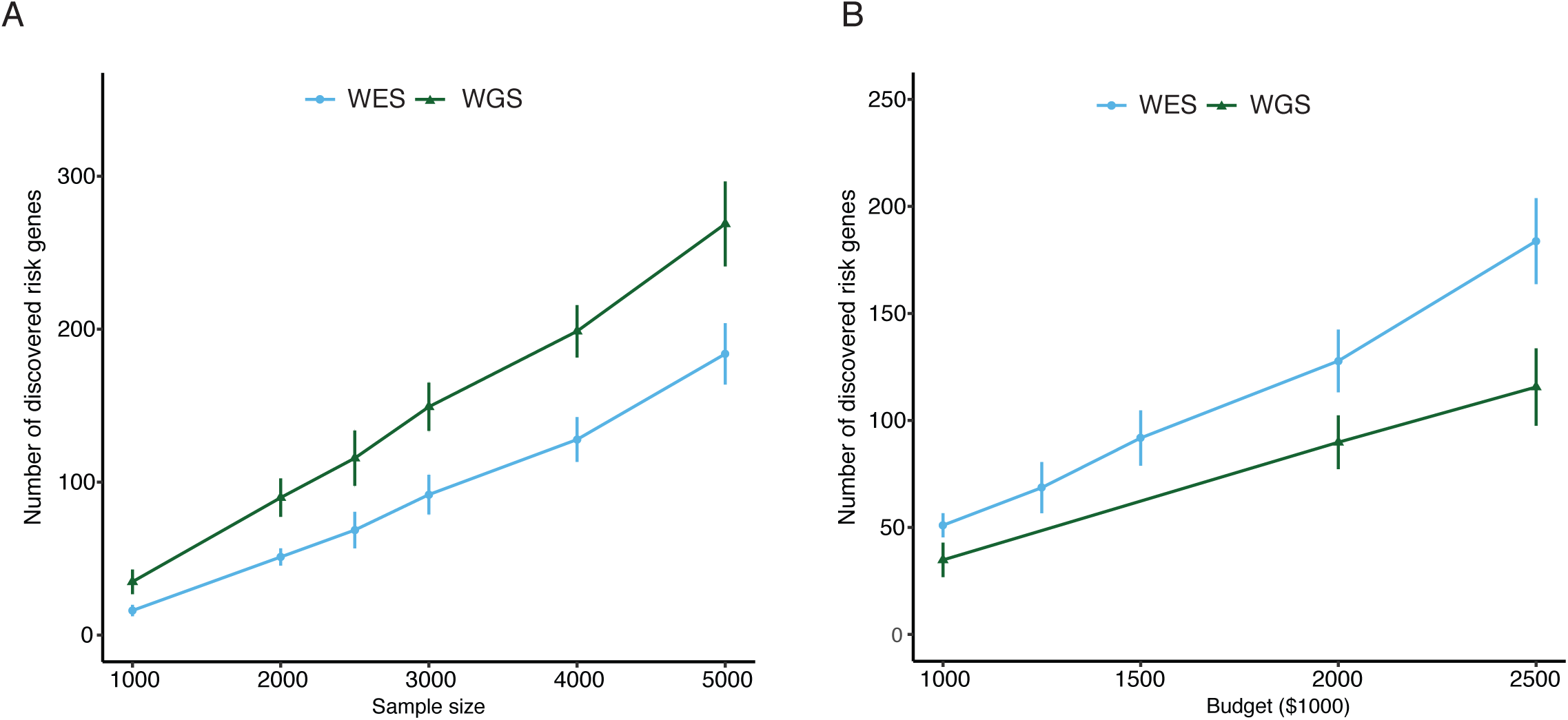
Comparison of power between WES and WGS from simulations. Power is measured as the number of discovered ASD risk genes at q-value < 0.1, and is obtained at each level of sample size (A) and sequencing cost (B).

## Discussion

Analyzing DNMs from exome sequencing data has been shown to be a powerful new paradigm for mapping risk genes of developmental and psychiatric disorders. Extending this to the non-coding genome is the natural next step and has the potential to transform our understanding of these complex disorders. In this work, we present a comprehensive statistical framework to support such analysis. The new method, TADA-A, is able to leverage multiple functional genomic annotations to better detect and prioritize risk-predisposing mutations. More importantly, TADA-A is able to combine information of all DNMs of a gene, in both coding and non-coding regions, to maximize the power to detect risk genes. The results of our meta-analysis of autism WGS datasets demonstrate the effectiveness of TADA-A. We show that *de novo* non-coding mutations make substantial contributions to the risk of autism (comparable to *de novo* LoF or missense mutations), and identified several promising new ASD risk genes. We note that the method can be applied to any units, other than genes, such as regulatory elements or sequence windows, though analysis at the gene level has the benefit of permitting us to borrow external information from previous WES studies.

A common strategy for analyzing DNM data is the burden analysis, which contrasts the rates of DNMs in affected individuals, often limited to likely functional mutations, with the expected rates due to chance alone. When researchers have no matched sibling or control data, the burden analysis can be confounded by technical factors such as sequencing depth. The burden analysis in non-coding regions is even more challenging because the statistical signal is considerably weaker than the coding signal, as reported by recent publications as well as our own study (Table 1). TADA-A greatly improves the standard burden analysis in several ways. Our mutation model, based on Poisson regression, incorporates covariates known to influence background mutation rates. In our ASD analysis, while we do not have access to genome-wide sequencing depth information, we used GC content as a proxy. Incorporating prior information of which genes are likely risk genes is critical for estimating parameters of annotations, while using a uniform prior largely lost the signals (Figure S2C).

One of the main challenges in making using of noncoding mutations in risk gene mapping is that we do not know a priori, from many possible noncoding annotations, which ones are disease relevant. TADA-A provides a convenient way to tackle this challenge. It allows users to analyze as many annotations as possible and learn which ones are informative of pathological mutations. In the application of TADA-A to ASD WGS data, we found that mutations with H3K27ac marks or with possible splicing effects contribute to ASD risk. These findings are consistent with previous research implicating a role of transcriptional mis-regulation in ASD etiology: chromatin remodeling and histone modification have been implicated in genes with ASD-associated DNMs;^1, 54^ trans-acting splicing modulators, such as FMRP, have been identified as syndromic ASD genes;^55^ and atypical splicing patterns of synaptic genes have been observed in individuals with autism.^56–58^ While we think enhancers (as marked by H3K27ac) and splicing regulations are involved in possibly most complex diseases, the exact annotations that are informative of disease variants may differ from our findings. For instance, it is possible that enhancers in only specific tissues, which are not known a priori, may be relevant to a given disease. And one may need to intersect H3K27ac with other annotations, e.g. conservation or open chromatin, to better identify functionally active enhancers. TADA-A provides an automatic way of learning such annotations (and their combinations).

Previous knowledge of the role of non-coding variants in diseases comes mostly from GWAS. The challenge with GWAS is that regulatory elements are much shorter (∼1kb) than regions of linkage disequilibrium (LD, hundreds of kb on average). It is thus not straightforward to assess the contribution of non-coding variants, or identify specific regulatory elements from GWAS.^45^ Indeed, the estimated contribution of DHS sites to heritability of complex diseases ranges widely from 79% to 25% in literature,^45, 59^ largely because of LD. By using DNMs, our work provides independent estimation of the contribution of both coding and non-coding variants to the risk of complex diseases.^45^ We estimated a modest average relative risk of about 1.5 for *de novo* mutations in less conserved brain H3K27ac enhancers and 3.4 in conserved brain H3K27ac enhancers, compared to 4-5 for missense and 20 for LoF mutations. We were not able to detect signal in evolutionarily conserved sequences (GERP and PhyloP, if not combined with tissue specific enhancers) or putative deleterious variants (CADD, trained mostly from non-brain tissues). These results suggest that evolutionary constraint is only weakly correlated with pathogenicity in ASD,^32^ and that regulatory variants of ASD probably act in a tissue and time-specific manner.^60^ Compared to a previous study,^47^ our estimate of ASD risk attributable to coding mutations is somewhat higher (1.9% vs. 1.1%), mainly due to a significant contribution from missense DNMs (0.83% vs. the previous estimate of 0.04%). We believe this difference is due to our different modeling assumptions: we treated all mutations in a category as a mixture of causal and non-causal mutations, whereas the previous study treated all mutations in a category equally (see Methods).^47^ We estimated that *de* novo coding (1.9%), non-coding (1.16%) and copy number variants (1.46%, estimated by a previous study^47^) together contribute 4.5% of ASD risk. We think that we significantly under-estimated the contribution of non-coding mutations to ASD risk for several reasons. First, we considered only enhancers within 10 kb of TSSs, which constitute about 36% of all enhancers in our data. Second, our dataset contains only regulatory sequences active early in development (5, 7, and 12 weeks post-conception) or in the adult brain. Third, larger genomic alterations, such as indels, potentially have larger effect sizes and are expected to increase the power for risk gene prediction. However, the false positive rates of calling *de novo* indels are much higher than SNVs. Besides, there is no good *de novo* mutation rate model for indels, which makes it difficult to model indels and estimate the relevant parameters. Thus *de novo* indels were not considered in this study. We also did not include *de novo* CNVs because of the difficulty of estimating mutation rates and attributing the contribution of a CNV to a risk gene.

Iossifov et al. used ascertainment differential, defined as the difference of DNM rates between probands and unaffected siblings, to measure the contribution of DNMs to ASD risk.^14^ Based on higher DN nonsynonymous mutation rates in probands, they estimate that DNMs contributes to about 21% cases. We note, however, this does not mean that the DNMs explain all these cases, since the DNMs are rarely fully penetrant. A better approach would estimate the contribution of DNMs to the disease liability, similar to the widely used heritability analysis, by taking into account the effect sizes of variants. Using this approach, both Gaugler et al. and our method reach similar estimates that DNMs contribute to a few percent of the ASD risk.^47^ To appreciate the difference of Iossifov et al. and the liability approach, consider a two-hit model where an individual has high ASD liability from inherited variants and one DNM with small effect pushing him above the liability threshold. In this case, DNM certainly contributes, but its effect is small. The ascertainment differential approach will not give us a correct picture of the true impact of DNMs in this scenario.

We identified four new ASD risk genes, three of which are strongly supported by other evidence. APBB1 is an adaptor protein localized in the nucleus. It is down-regulated in ASD cerebellum than control,^61^ and its microexons are mis-regulated in the brains of ASD individuals.^56^ NRXN1 belongs to a group of presynaptic cell adhesion molecules which controls synapse development.^62^ It has been implicated as a top candidate gene for neurodevelopmental and neuropsychiatric conditions.^63^ Interestingly, alternative splicing of *Nrxn1* has been reported to cause defects of synaptic formation in the hippocampus region in a mouse model (the gene is supported by a splicing SNV, Table 2).^64^ TANC2 is a member of postsynaptic scaffold proteins. It is highly expressed in the brain and play roles in the regulation of dendritic spines and excitatory synapses.^65^ One WES study reported TANC2 as a candidate intellectual disability gene.^66^

Most of the new ASD genes at FDR < 0.3 are supported by functional or association studies (summarized in Table S6). *JUP* is a member of the catenin/cadherin superfamily which has important roles in neuron connections and interactions. ^67^It is strongly expressed in the primate prefrontal cortex and hippocampus.^68^ *Dll1* is expressed in most of the neural tube during CNS development in mice.^69^ Studies of Dll1 deficient mice suggest that Dll1 play an important role in the expansion and differentiation of mesencephalic dopaminergic neural precursor cells into neurons.^70^ *PPM1D* has recently been identified as a novel risk gene for intellectual disability.^71^ *MSL2*, *DLL1, SMARCC2, ARHGAP44* and *GAPVD1* are predicted as autism risk genes by a recently developed machine learning approach that utilizes a brain-specific functional interaction network^24^ (q-values 0.038, 0.0186, 0.0319, 0.0859 and 0.08 respectively).

We also found that the two TAD regions with excess regulatory SNVs in ASD are supported by CNV studies. In one TAD, recurrent, rare CNVs (Chr2: 45455651-45984915) spanning the entire *SRBD1* gene (the only protein coding gene disrupted by the CNVs within this TAD) were reported in ASD subjects.^72^ In a later independent study, CNVs in this TAD region were found to be enriched in ASD cases versus controls.^72^ These results suggest that *SRBD1* is likely the risk gene in this TAD. In the other TAD region, ASD-associated duplication of 8p23.1-8p23.2 introduces a breakpoint between MSRA and RP1L1.^73^ MSRA is a member of the methionine-sulfoxide reductase system whose function is to alleviate oxidative stress. Increased exposure to oxidative stress plays an important role in the pathogenesis of ASD.^74^ In addition, GWAS studies have established associations of MSRA with schizophrenia^75^ and bipolar disorder.^76^

One caveat of using multiple dataset for meta-analysis using TADA-A is that DNM load is likely to be different between simplex and multiplex families. Ideally, we would like to treat the data from simplex and multiplex families differently, but in practice, this would reduce sample size and make the estimates less reliable. Several lines of reasoning/evidence suggest that the extent of difference may be limited. (1) When choosing simplex families, it is hard to exclude families with high genetic risks, because the family sizes are often small. A high risk family by chance could give rise to two affected siblings or one affected and one unaffected siblings. The former would be classified as multiplex and the latter simplex. It is estimated that more than 85% of such high risk families with two children, at least one with autism, would be included in Simons Simplex Collection.^77^ (2) The *de novo* CNV burden of simplex and multiple families were found to be similar. In Pinto et al, the rates of de novo CNVs is 5.9% in simplex and 5.8% in multiplex families.^78^ (3) The burden of nonsynonymous mutations from our data (Figure 1B, OR about 1.2), in which multiplex families from Yuen et al. takes a majority proportion, is quite similar to the burden based on simplex families.^14^ (4) Yuen et al. found identical DNMs existed in 19% of the sibling pairs of multiplex families they investigated.^79^ This observation suggests that, even in multiplex families, DNMs derived from germ line mosaic mutations could play a significant role in increasing ASD risk. These “mosaic” DNMs are thus similar to DNMs in simplex families, in a sense. With more data from simplex and multiplex families available, accounting for this difference would be a future direction.

We believe that TADA-A can be further developed along several directions. The baseline mutation model of TADA-A is relatively simple, and recent studies demonstrate that broader sequence context and additional genomic features can be highly correlated with mutation rates.^80^ Additionally, in the current analysis, we focus on regulatory sequences close to genes. However, a large fraction of regulatory sequences are distal to transcription start sites. The challenge is that the target genes of these sequences are often unknown. We plan to integrate chromatin interaction data (e.g. Hi-C) in the future to better analyze mutations in distal enhancers. Finally, TADA-A uses a linear model for predicting effects of mutations from annotations. A more powerful method may use a non-linear model such as deep neuron networks.^81^

## Acknowledgements

This work was supported by National Institutes of Health grant (1R01MH110531) and Simons Foundation award (SFARI Award ID 385027) to X.H.

## Web Resources

Wu et al. WGS data: http://wwwdev.ebi.ac.uk/eva/?eva-study=PRJEB14713

TADA-A: https://github.com/TADA-A/TADA-A

CADD: http://cadd.gs.washington.edu/score

SFARI genes: https://sfari.org

BrainSpan: http://developinghumanbrain.org

Cross-tissue enhancer/promoter correlation: http://khuranalab.med.cornell.edu/roadmap_stringent_enhancers.txt

## Supplemental Data

Supplemental data include 4 figures and 8 tables.

Figure S1. Distributions of the distances between splicing mutations to their nearest exon/intron junctions. (A), (B), (C), and (D) represent mutations with mutant allele as “A”, “T”, “C”, and “G”, respectively.

Figure S2. Relative risk estimates of annotations using different priors for ASD data.

Figure S3. TADs enriched for regulatory SNVs. The heatmaps represent the interaction strength between two genomic loci measured from Hi-C experiments.

Figure S4. Relative risk estimates of annotations for ASD data after filtering out mutations with allele frequency greater than 0.01 in GnomAD or BRAVO.

Table S1. Summary of studies contributing DNMs.

Table S2. Calibration of mutation rates for each study.

Table S3. Joint estimates of the relative risks of noncoding annotations that passed feature selection.

Table S4. Mutational counts and evidence of 12 new ASD genes (FDR < 0.3).

Table S5. Details of geneset enrichment analysis.

Table S6. Functional relevance and literature support of new ASD genes.

Table S7. Enhancers with recurrent *de novo* SNVs. Genes were assigned to enhancers based on cross-tissue correlation between enhancer activity and gene expression from Roadmap Epigenomics (See Web Resources).

Table S8. TAD-level analysis for SNVs. The table show the top two TADs based on Poisson test (comparing observed and expected number of mutations).

