## Supplementary Materials for "A statistical framework for mapping risk genes from *de novo* mutations in whole-genome sequencing studies"

Figure S1

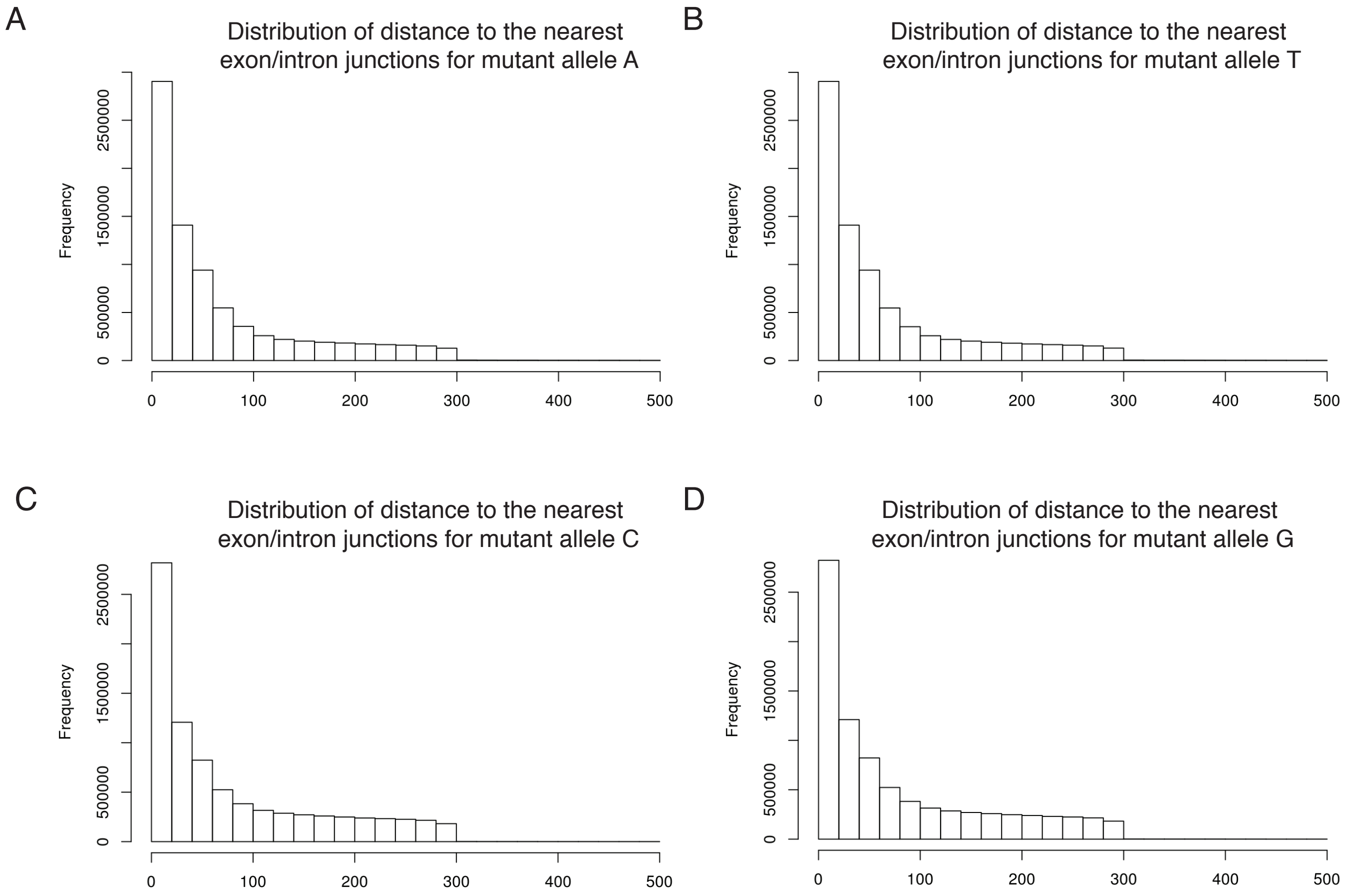


Figure S1. Distributions of the distances between splicing mutations to their nearest exon/intron junctions. (A), (B), (C), and (D) represent mutations with mutant allele as "A", "T", "C", and "G", respectively.

Figure S2

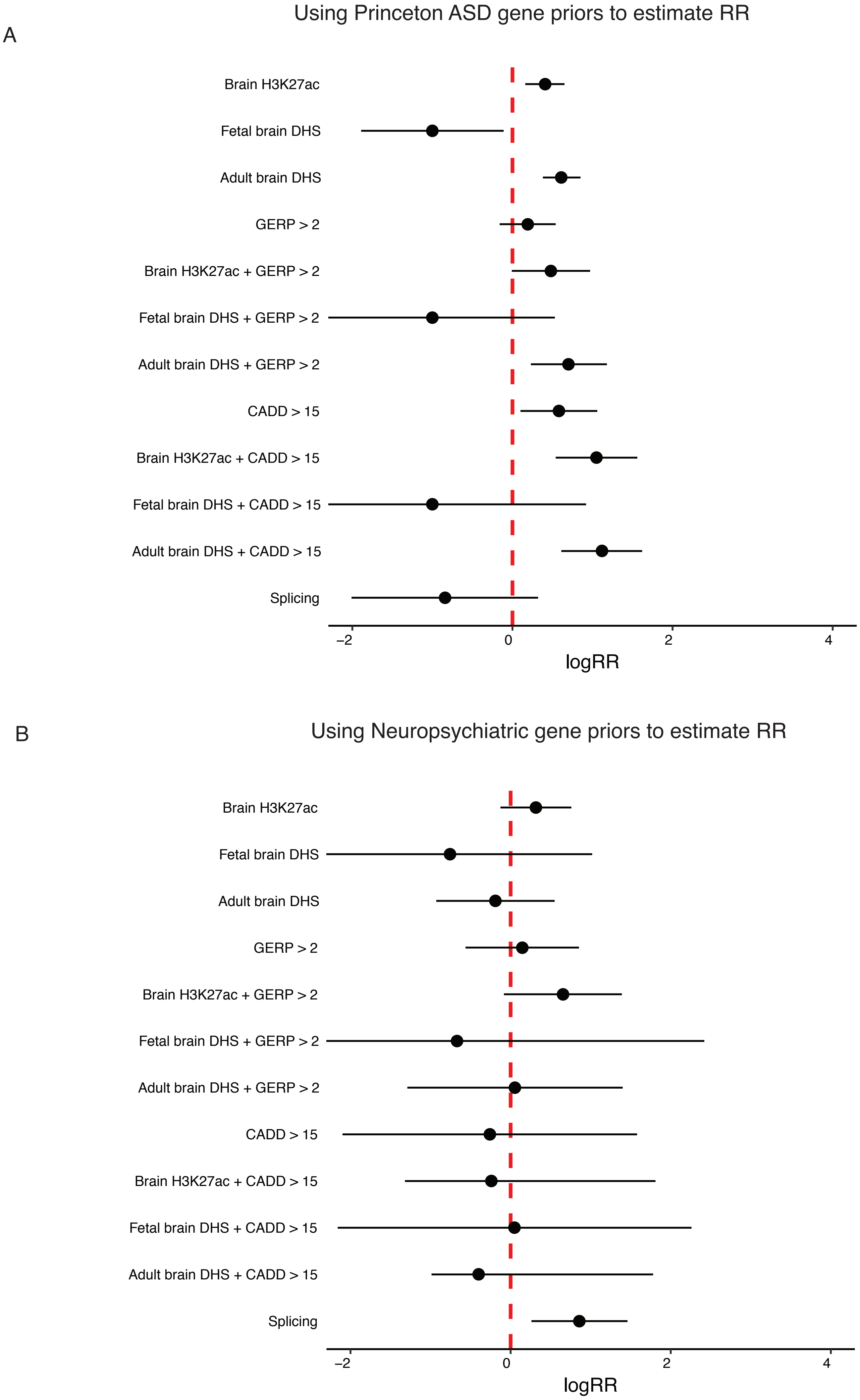
A
Using Princeton ASD gene priors to estimate RR
Brain H3K27ac
Fetal brain DHS
Adult brain DHS
GERP > 2
Brain H3K27ac + GERP > 2
Fetal brain DHS + GERP > 2
Adult brain DHS + GERP > 2
CADD > 15
Brain H3K27ac + CADD > 15
Fetal brain DHS + CADD > 15
Adult brain DHS + CADD > 15
Splicing
−2
0
2
4
logRR
B
Using Neuropsychiatric gene priors to estimate RR
Brain H3K27ac
Fetal brain DHS
Adult brain DHS
GERP > 2
Brain H3K27ac + GERP > 2
Fetal brain DHS + GERP > 2
Adult brain DHS + GERP > 2
CADD > 15
Brain H3K27ac + CADD > 15
Fetal brain DHS + CADD > 15
Adult brain DHS + CADD > 15
Splicing
−2
0
2
4
logRR

Figure S2

C

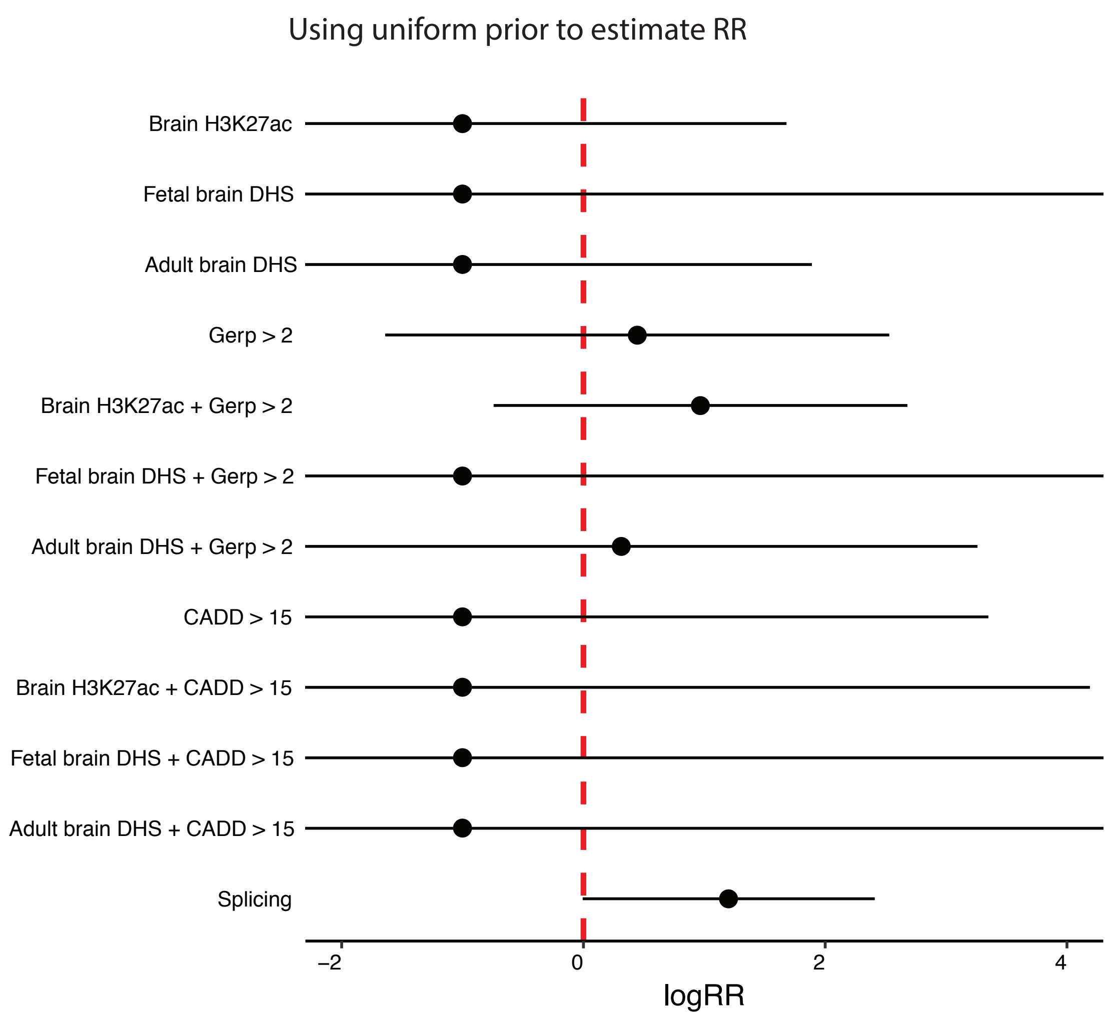


Figure S2. Relative risk estimates of annotations using different priors for ASD data.

Figure S3

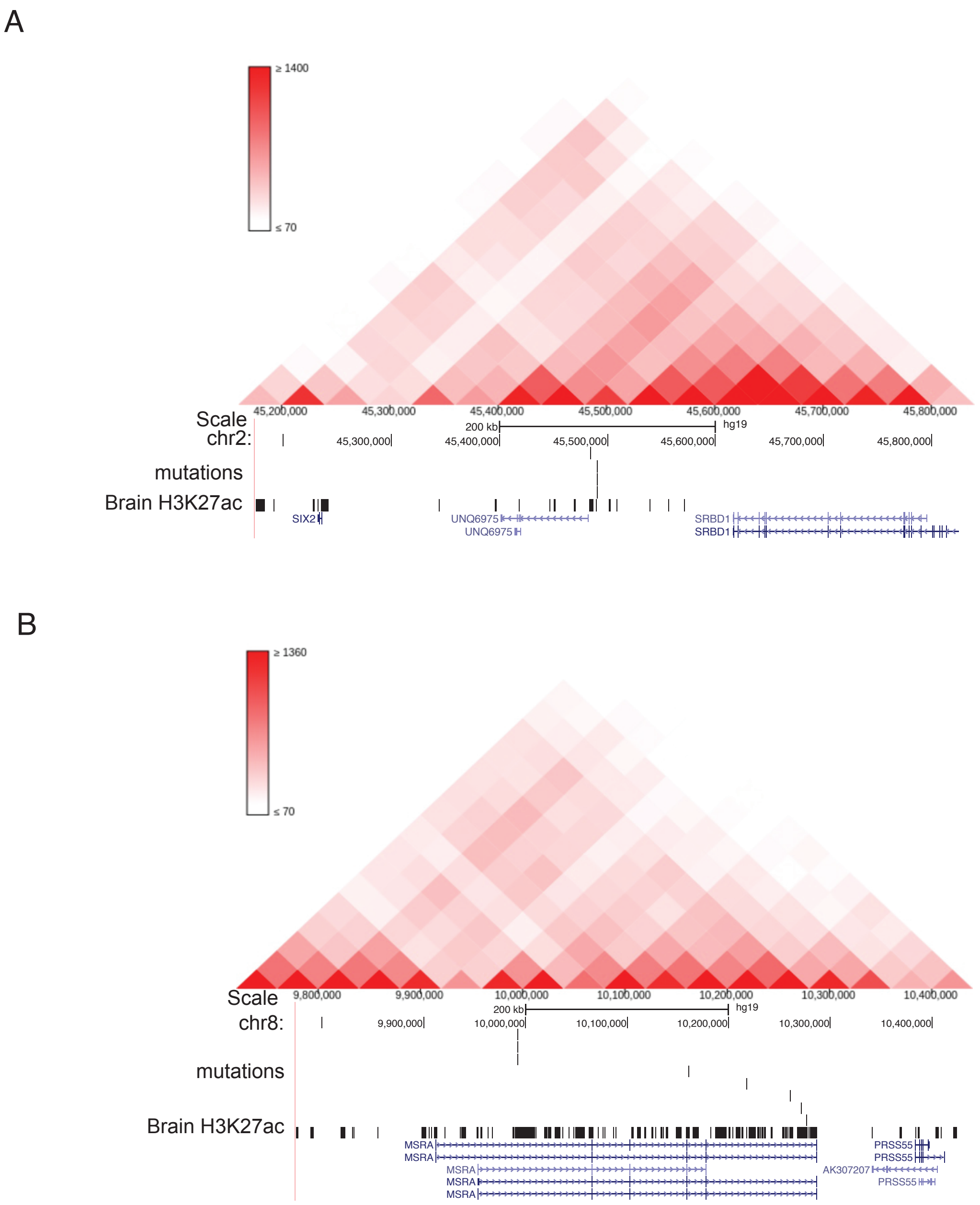


Figure S3. TADs enriched for regulatory SNVs. The heatmaps represent the interaction strength between two genomic loci measured from Hi-C experiments.

Figure S4

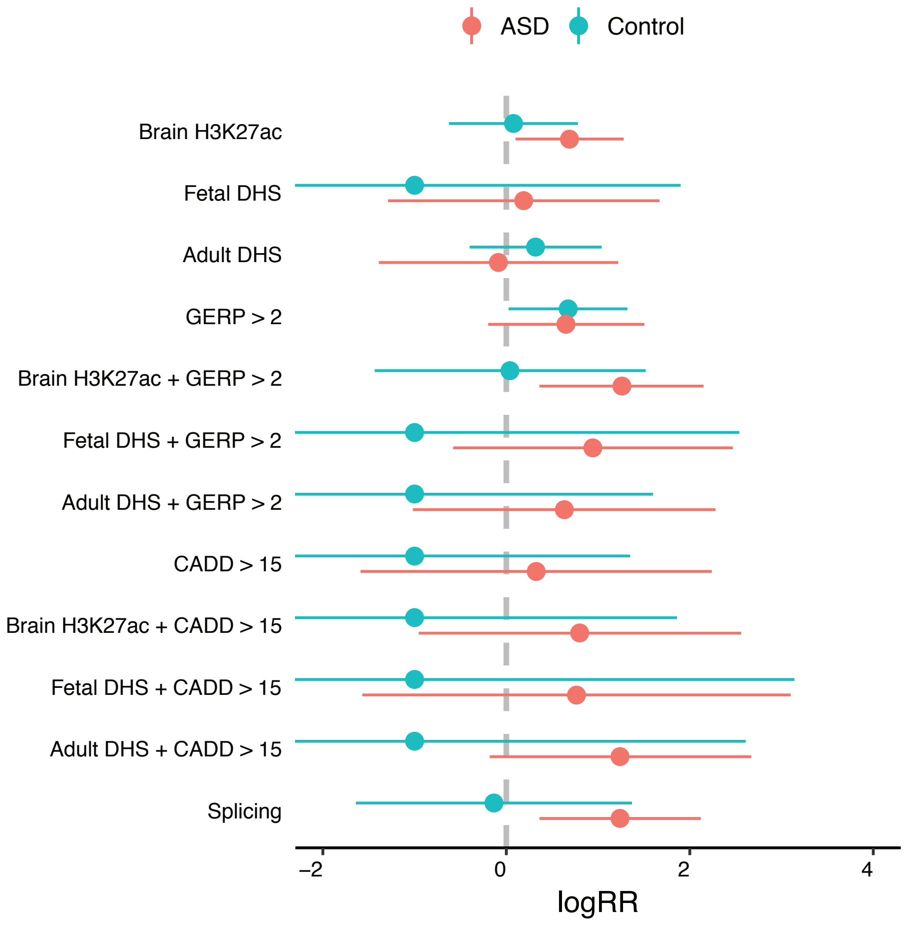


Figure S4. Relative risk estimates of annotations for ASD data after filtering out mutations with allele frequency greater than 0.01 in GnomAD or BRAVO.

### Supplemental Methods

#### 1 TADA-A: computational speedup for parameter estimation

Fitting the TADA-A model at the base level can be very computationally expensive: even if we apply our model only to 10kb regions near TSS, we have about 200M positions. We describe our computational strategy to speed up the algorithm. We note that the idea can be applied to both the mutation and functional model of TADA-A because of similar mathematical structure. For now, we consider our model of mutation counts of a gene assuming it is a risk gene. Let $y_{it}, \mu_{it}$ be the mutation count and calibrated mutation rate of position $i$ mutating to allele $t$, respectively. Suppose there are $K$ annotations, and define $x_{it}$ as the vector of annotations of $i_t$, the mutant allele $t$ at position $i$ ($K$-dimension). Our model is:

$$y_{it} \sim \text{Poisson}(\mu_{it} e^{x_{it}\beta}), \tag{1}$$

where $\beta$ is the effect of annotations ($K$-dimension vector). Note that in this equation, we drop the constant $2N$ in mutation rates for simplicity. So $\mu_{it}$ should be interpreted as the expected number of mutations of allele $t$ in position $i$ (rather than rate per chromosome). For simplicity, we assume the $k$-th annotation of $i_t$, $x_{itk}$, is binary. Our goal is to quickly evaluate the likelihood of all bases in a gene. It is easy to show that the log-likelihood as:

$$\log P(y|\beta) = \sum_t (\sum_i y_{it} \log \mu_{it} + \sum_i y_{it}(x_{it}\beta) - \sum_i \mu_{it} e^{x_{it}\beta} - \sum_i \log y_{it}!\,). \tag{2}$$

The terms $\sum_i y_{it} \log \mu_{it}$, $\sum_i \log y_{it}!$ do not depend on annotations and can be easy to deal with (we will show below that these constant terms cancel out in the final likelihood). For the other two terms, we take advantage of this simple fact: all position/mutant-allele pairs with identical annotations would have the same value of $x_{it}\beta$ and $e^{x_{it}\beta}$. This greatly simplifies the log-likelihood. We consider all possible combinations of annotations, and let $c$ be a category representing one combination ($2^K$ possible categories in theory). Ex. a category might be: conserved, open chromatin in brain, but not motif, represented as (1,1,0) for the three annotations: conservation, chromatin accessibility and motif. We define $V_{ck}$ be an indicator variable of whether the $k$-annotation is 1 under the category $c$. We define the variable $\gamma_c$:

$$\gamma_c(\beta) = \exp\left(\sum_k \beta_k V_{ck}\right). \tag{3}$$

This can be interpreted as the relative risk of mutations belonging to the category $c$. Now for a mutant of allele $t$ at position $i$ belonging to category $c$, the terms $x_{it}\beta$ and $e^{x_{it}\beta}$ become $\log(\gamma_c)$ and $\gamma_c$, respectively. The log-likelihood of a gene is now written as:

$$\log P(y|\beta) = \sum_c y_c \log(\gamma_c) - \sum_c \mu_c \gamma_c + \sum_t (\sum_i y_{it} \log \mu_{it} - \sum_i \log y_{it}!\,). \tag{4}$$

where $y_c$ and $\mu_c$ are the total DNM count and total mutation rates of category $c$ respectively. Computation of this function is many times faster than the naive implementation because we do not have to compute $x_{it}\beta$ and $e^{x_{it}\beta}$ for all positions.

If a gene is a non-risk gene, its likelihood is easy to evaluate. For each base $i$ mutating to allele $t$, we have $y_{it} \sim \text{Poisson}(\mu_{it})$. This leads to:

$$\log P(y|\beta) = \sum_t (\sum_i y_{it} \log \mu_{it} - \sum_i \log y_{it}!\,) - \mu, \tag{5}$$

where $\mu$ is the total mutation rate of the gene, summing over all the positions and all possible mutant alleles.

Now we consider the total likelihood over all genes, defined in the main text as:

$$P(Y|\beta, \pi) = \prod_g [\pi_g P(Y_g|Z_g = 1, \beta) + (1 - \pi_g) P(Y_g|Z_g = 0)] \quad (6)$$

We can factorize the term $P(Y_g|Z_g = 0)$, which does not depend on the parameters. Let $B_g$ be the Bayes factor of gene $g$, $B_g = P(Y_g|Z_g = 1, \beta)/P(Y_g|Z_g = 0)$, the likelihood can be written as:

$$P(Y|\beta, \pi) \propto \prod_g [\pi_g B_g(\beta) + (1 - \pi_g)] \quad (7)$$

Using Equations 4 and 5, we obtain the log BF of a gene as:

$$\log B_g(\beta) = \sum_c y_c \log(\gamma_c(\beta)) - \sum_c \mu_c \gamma_c(\beta) + \mu. \quad (8)$$

We note that the idea of categorization is not new: it underlies the original TADA model. The difference here is that: we use categorization only for computational purpose, and our statistical model does not make this assumption. Had we applied the original TADA directly here, we will have $2^K$ parameters (relative risk for each category), while we only have $K$ parameters under TADA-A. When the annotations are continuous, we could do more refined discretization, but the number of parameters would be even higher under the original TADA, but does not change under TADA-A.

#### 2 TADA-A: additional details

The BF of a gene has a simple interpretation. We subtract Equations 5 from 2, and obtain the log-BF of a gene as:

$$\log B = \sum_t (\sum_i y_{it} \log(\gamma_{it})) - \sum_t (\sum_i \mu_{it}(\gamma_{it} - 1)), \quad (9)$$

where $\gamma_{it}$ is the relative risk of mutation of allele $t$ at position $i$. We assume there are no protective mutations, i.e. $\gamma_{it} \geq 1$. The first term is the weighted mutation count of the gene, where weight is given by $\log \gamma_{it}$ (non-negative). More damaging mutations would thus contribute more to the positive score than less damaging ones. The second, negative term represents the penalty given to the gene. The penalty is larger for larger genes, genes with high mutation rates, and with more positions predicted to be damaging (large $\gamma_{it}$ means we expect more DNMs under the risk gene model, so if we do not observe DNM of allele $t$ at $i$, we have more penalty).

#### 3 Calling *de novo* SNVs from 32 new ASD trios

32 unrelated ASD patients of Han Chinese ancestry (30 males and 2 females) and their unaffected parents were recruited for this study. Diagnostic and Statistical Manual of Mental Disorders-4th edition (DSM-IV-TR) and Autism Diagnostic Observation Schedule (ADOS) were employed by autism specialists for diagnosis. Genomic DNAs from 96 individuals were used to construct genomic DNA libraries (500-bp), followed by Illumina paired-end sequencing (90-bp).

Trim Galore (`http://www.bioinformatics.babraham.ac.uk/projects/trim_galore/`) and Cutadapt [1] were used to remove $3'/5'$ adapters and low quality reads from raw data. The pre-processed reads were aligned to the human reference genome (hg19, GRCh37) by Burrows-Wheeler

Aligner (BWA) [2] allowing at most four mismatches. An average of 108 Gb of reads per individual were generated with 36-fold coverage. Then, Sequence Alignment/Map tools (SAMtools) [3] was used to mark and remove duplicate reads. More than 99% and 98% of the human genome is covered by at least one and at least 10 reads, respectively. We applied the Genome Analysis Toolkit (GATK) [4] on uniquely mapped reads to detect single nucleotide variants (SNVs). Variants were detected on all three individuals in each trio simultaneously.

We used ForestDNM to call de novo SNVs as described previously in [5][6]. The method used in Jiang et al. (2013) was employed to call de novo indels. Since DNMs are very rare, we removed any variants with a minor allele frequency ($MAF$) $> 0.1\%$ based on allele frequency from dbSNP138 and 1000 Genomes to improve the accuracy of DNM calling. The total number of DNMs we called were similar to previous reports using the same methods [5][6].

#### 4 Contributions of various mutational categories to autism risk

We consider multiple types of coding and noncoding de novo mutations, and estimate how much they contribute to the risk of autism, based on a liability model. Without loss of generality, we assumed ASD liability in the general population follows a standard normal distribution $Y \sim N(0, 1)$. Individuals carrying mutations have a shifted liability distribution

$$Y \sim N\left(\sum_j \beta_j X_j, 1\right) \tag{10}$$

where $\beta_j$ is the effect size of the $j$-th mutation type at the liability scale, $X_j$ is an indicator variable representing if an individual has the $j$-th mutation type or not. Let $p_j$ be the probability of having one $j$-th type mutation in any ASD risk gene, and $\pi$ be the proportion of ASD risk genes. Given that mutation occurs independently, the variance of liabaility due to all mutations is:

$$V = \sum_j \beta_j^2 \mathrm{Var}\left(X_j\right) = \sum_j \beta_j^2 p_j (1 - p_j) \tag{11}$$

with the variance explained by type $j$ mutation:

$$V_j = \beta_j^2 p_j (1 - p_j) \tag{12}$$

The mutation rate for a type $j$ above can be simply obtained by multiplying the total mutation rate of type $j$ and $\pi$. We chose $\pi = 0.06$ in this study, based on several independent studies [7][8].

The relative risks of coding LoF mutations and Mis3 mutations are derived from a previous ASD WES study, as 20 and 4.7, respectively. We used WGS data to derive the relative risks of non-coding mutations (Table 1). The relative risks of less conserved regulatory SNVs, conserved regulatory SNVs and splicing SNVs are 1.55, 3.46 and 3.27, respectively.

We assume each affected individual only has one type of mutation from risk genes, which is a reasonable assumption given the DNM rate is low. Let $s_0$ be the prevalence of ASD in the general population (use $1/68 = 0.0147$), and $s_j$ be the proportion of ASD patients among all with $j-$th type mutation. By definition of relative risk, $\bar{\gamma}^{(j)} = s_j / s_0$. It is straight forward to relate effect size at liability scale and relative risk:

$$\beta_j = \phi^{-1}(1 - s_0) - \phi^{(-1)}(1 - s_j) = \phi^{-1}(1 - s_0) - \phi^{(-1)}(1 - s_0 \bar{\gamma}^{(j)}) \tag{13}$$

where $\phi^{-1}$is the inverse cumulative distribution function of a standard normal distribution. Plug in $p_j$ and $\beta_j$ to Equation 12, we can obtain the variance of risk explaiend by each type of mutation. In the paper, we also report the relative proportion of de novo ASD risk explained by any type of mutation, $V_j/V$.

#### 5 Simulation studies to compare the power of WES and WGS

We used simulations to generate coding and non-coding DNMs from ASD risk genes and non-risk genes and then used TADA-A to call ASD risk genes, either by using only coding mutations (WES approach)or using both coding and non-coding mutations (WGS approach). We performed 5 simulations at each different number of $N$ trios. For each iteration at each value of $N$, we run the following steps:

1. We randomly chose ASD risk genes from 18700 genes based on a binomial distribution with success probability of 0.06, and the rest of genes are ASD non-risk genes.

2. For each gene, we generated mutations of each possible allele $t$ at each position $i$ based on on $Poisson$ distribution (Equation 1). For risk genes, the estimates of $exp(\beta)$ is in Table 1 in the main text. For non-risk genes, $\beta$ is set to be 0.

3. We used TADA-A to identify ASD risk genes at $q < 0.1$, by using both coding mutations (WES approach) or by using both coding and non-coding mutations (WGS approach).

4. We identified the number of true ASD risk genes (defined in Step 1) that have been identified in the last step by WES and WGS approaches, respectively.
